# Arginine metabolism has a pivotal function for the encystation of *Giardia duodenalis*

**DOI:** 10.1101/2025.02.20.639220

**Authors:** Christian Klotz, Ricarda Leisering, Kari D Hagen, Hannah N Starcevich, Christoph Ewald, Samuel Türken, Malte Marquardt, Saskia Schramm, Totta Ehret Kasemo, Stefanie Marek, Frank Seeber, Ralf Ignatius, Scott Dawson, Toni Aebischer

## Abstract

Arginine metabolism plays a key role in the energy metabolism of the intestinal parasite *Giardia duodenalis*, an amitochondrial protozoan that infects humans and animals and causes significant morbidity. An arginine deiminase (ADI) has been implicated in virulence, but it is currently unknown if ADI allele variants from the different genetic *G. duodenalis* subgroups (assemblages) differ in function. Here, the hypothesis was tested that sequence variation detected between *G. duodenalis* ADI alleles from the two *G. duodenalis* assemblage types found in humans affects functional parameters of the enzyme with potential consequences in life cycle progression.

The ADI enzyme affinity for arginine was drastically reduced in sub-assemblage AII isolates, a human specific assemblage, in comparison to zoonotic sub-assemblage AI and B isolates. We identified the two amino acid residues responsible for the lower substrate affinity of ADI_AII_ variant. Using genetic approaches to generate ADI knockout mutants, biochemical approaches to unravel substrate affinity as well as cellular approaches to determine efficiency of life cycle progression, we show that ADI is essential for efficient encystation of the parasite and that the lower substrate affinity in ADI_AII_ correlates with lower encystation efficiency. We further demonstrate that arginine is essential for efficient encystation, and by generating ADI knock-out parasites we present evidence that ADI is the functional correlate for this arginine dependence.

Thus, our data describe ADI as a quantitative trait that affects life cycle progression of *G. duodenalis* with putative clinical and epidemiological relevance.

**Author summary:** In the human pathogenic parasite *Giardia duodenalis*, arginine deiminase (ADI) mediates the first step in the arginine dehydrolase pathway (ADH), metabolizing arginine to provide chemical energy in form of ATP. The bacterial-derived ADH pathway had been inherited by horizontal gene transfer, and ADI has been proposed as a virulence factor. We show here by biochemical and genetic approaches with ADI knock-out mutants that arginine and its metabolizing enzyme ADI are essential for efficient life cycle progression (encystation) to form infectious cysts. Furthermore, we show a drastically impaired arginine substrate affinity for the human-specific *G. duodenalis* genotype AII in comparison to the zoonotic genotypes AI and B and identified the molecular entities responsible for this altered substrate affinity. This lower substrate affinity also correlated with lower cyst formation in the AII genotype.

## Introduction

*Giardia duodenalis* (syn. *G. lamblia*, *G. intestinalis*) is a medically important protozoan parasite infecting the small intestine of mammalian hosts. Advances in genotyping and its application to epidemiological studies revealed a complex population structure [1, 2]. According to genetic information, these parasites can be grouped into eight distinct assemblages (A-H) [1]. Of these, assemblages A and B are found in humans and a broad range of domestic and wildlife animals [1]. The remaining assemblages show more narrow host ranges, with assemblage C and D found in canides, E in hoofed animals, F in felids and H in mammalian sea life [1].

Infections with different assemblages have been correlated with differences in clinical symptoms [3], and although this remains a matter of debate, it is implying that differences in virulence between parasites are genetically encoded. Interestingly, the proportions of sporadic assemblage A and assemblage B infections in humans - which are not associated with outbreaks - are uneven, with assemblage B being overall more prevalent [3–6]. Recently, improved genetic typing schemes revealed that human assemblage A infections are predominantly caused by sub-assemblage AII, a sub-assemblage type only found in humans, and only rarely by assemblage AI, a sub-assemblage of animals and humans [6–8]. The molecular mechanisms as well as involved parasite products that lead to clinical disease and may affect virulence as well as underlay differences in assemblage distribution in humans remain ill-defined [9–12]. However, several molecular virulence factors have been proposed [9, 10]. Arginine deiminase (ADI) is one of these [13–15] and is thought to provide multiple virulence-associated functions in the parasite and its interaction with the host. For example, *G. duodenalis* ADI has been shown to potentially interfere with NO-dependent anti-parasite defense [16, 17] and also with epithelial cell proliferation through arginine consumption [18]. Competition by parasites for arginine with the host epithelium or microbiome thus may induce host pathobiology and immune modulation [16, 17].

The ADI enzyme in *G. duodenalis* mediates the first catalytic step of the arginine dihydrolase (ADH) pathway (Fig 1A) and catalyzes the conversion of arginine to citrulline releasing NH_3_ [19]. Notably, ADI of *G. duodenalis* was an early example of a trans-kingdom horizontal gene transfer from pro- to eukaryotes [19].

**Figure 1.**
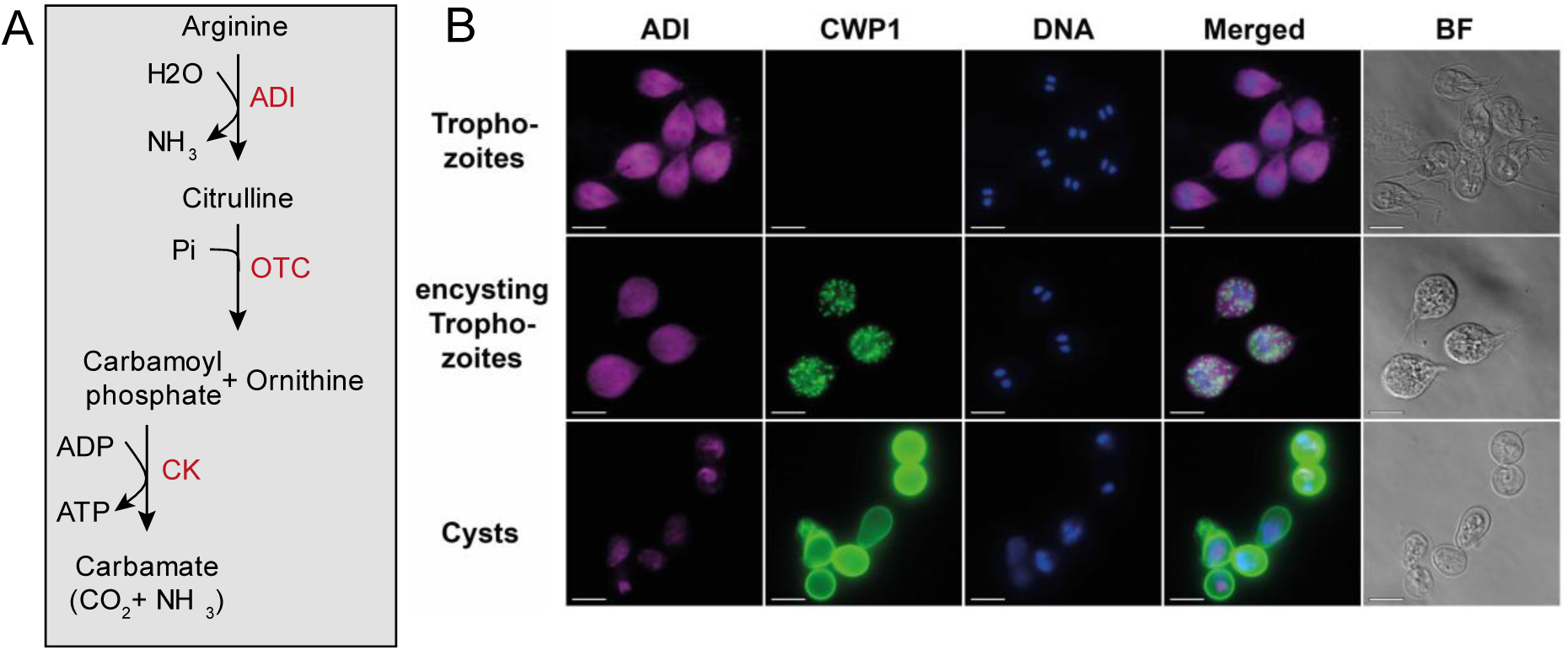
Sketch of the arginine dehydrolase (ADH) pathway and localization of arginine deiminase (ADI) in different life cycle stages of *G. duodenlis*. (A) Overview of the ATP-generating ADH-pathway, including the key enzymes in red: arginine deiminase (ADI), ornithine-transcarbamylase (OTC) and carbamate kinase (CK). (B) Cytoplasmatic localization of ADI in trophozoites and in vitro encysting trophozoites and cysts (the latter both 24 h after induction of encystation). Bright field (BF) and immunofluorescence images are shown using a polyclonal alpaca antiserum raised against recombinant ADI protein (magenta), anti CWP1-antibody, and nucleic acid stain Hoechst (blue). Scale bar = 10 µm.

As an amitochondrial parasite lacking respiratory metabolism, substrate level phosphorylation by the glycolytic, pyruvate, and the ADH pathways provides ATP and reducing equivalents for metabolism. While ATP generation by any of the three pathways may contribute to different aspects in *Giardia* biology, including life cycle transitions or metabolic interactions with the host, energy generation may also be limited or regulated by local substrate concentrations of glucose or arginine or local oxygen concentrations. Despite the lower energy yield, *Giardia* trophozoites prioritize ATP generation from arginine using the ADH pathway consisting of three enzymes (one net ATP, see Figure 1A) over energy production from glycolysis (two net ATPs). Enzymes of the ADH pathway are amongst the most highly expressed metabolic enzymes in trophozoites [20–23], which furthermore mirrors their importance.

We have previously shown that in vitro depletion of arginine by *G. duodenalis* ADI modulates cytokine production and surface marker expression of activated human monocyte-derived dendritic cells, with potential consequences for adaptive immune responses [24]. Thus, *G. duodenalis* ADI may influence parasite virulence, pathogenesis, and host immunity by several modes of action. *G. duodenalis* ADI has also been proposed to have a role during encystation through histone modification [15]. However, whether it really contributes to life cycle progression and, hence, transmission has not been further evaluated.

Comparison of ADI sequence information in genomes of the human relevant *G. duodenalis* (sub)-assemblages A and B [25–29] indicates various grades of sequence polymorphisms, in particular between assemblage A and B whose genomes only share approximately 70% sequence homology [27, 28, 30]. The ADI sequence is encoded by a single gene that contains only one exon encoding a protein of 580 amino acids. Whether the sequence polymorphisms of the ADI gene are functionally relevant has not been addressed thus far. However, this seems highly relevant given the mentioned roles of ADI in the biology of the parasite. Here, we show that two specific amino acid polymorphisms that distinguish ADI from assemblage AII from that in AI and B parasites are responsible for changes in the *K*_m_ values of the enzymes. They lead to lower substrate affinity of ADI_AII_. By knock-out experiments and reverse genetics we further show that the changes in biochemical characteristics translate to a lower encystation rate and thus have direct functional consequence. This provides the first evidence for a functional polymorphism of ADI as a virulence factor within the diverse *G. duodenalis* population.

## Results

The ADH pathway is implicated in key roles for microaerophilic *Giardia*, and as shown in Figure 1A, ADI catalyzes the first step in the ADH pathway by converting arginine into citrulline and ammonia. As seen in Figure 1B, immunofluorescence analysis revealed that ADI is localized to the cytoplasm of both metabolically active and encysting trophozoites. In the metabolically less active cyst form, however, the ADI signal was reduced and, with respect to localization, was unevenly distributed throughout the cytoplasm (Figure 1B). Both observations confirm previous reports of ADI protein abundance and localization in both life cycle stages and during encystation [22, 31].

### ADI allelic variants from human-derived G. duodenalis assemblage AII isolates possess lower arginine substrate affinity than ADI from zoonotic assemblages AI and B

Cellular energy metabolism, including arginine metabolism, may vary between different *G. duodenalis* assemblages. While ADI is known to play roles in diverse *G. duodenalis* virulence aspects, such as encystation [15], the relevance of putative ADI activity variants in distinct *G. duodenalis* assemblages and their role in virulence remains unknown and was therefore tested next.

To define potential functional variation of arginine metabolism in different *G. duodenalis* assemblages, we first investigated ADI sequence variation (Figure 2) and enzymatic activity (Figure 3A, B) in representative clinical *G. duodenalis* isolate cultures previously established in our laboratory. These included assemblage AII (isolates P407 and P506) and assemblage B (isolates P424 and P387) [28, 32] as compared to the well-studied reference strains WB (assemblage AI) and GS (assemblage B). Together, these isolates provide examples of assemblages AI, AII and B found in humans and are all culturable *in vitro*. Except for a few assemblage E isolates, no successful *in vitro* cultures of other *G. duodenalis* assemblages have been described.

**Figure 2.**
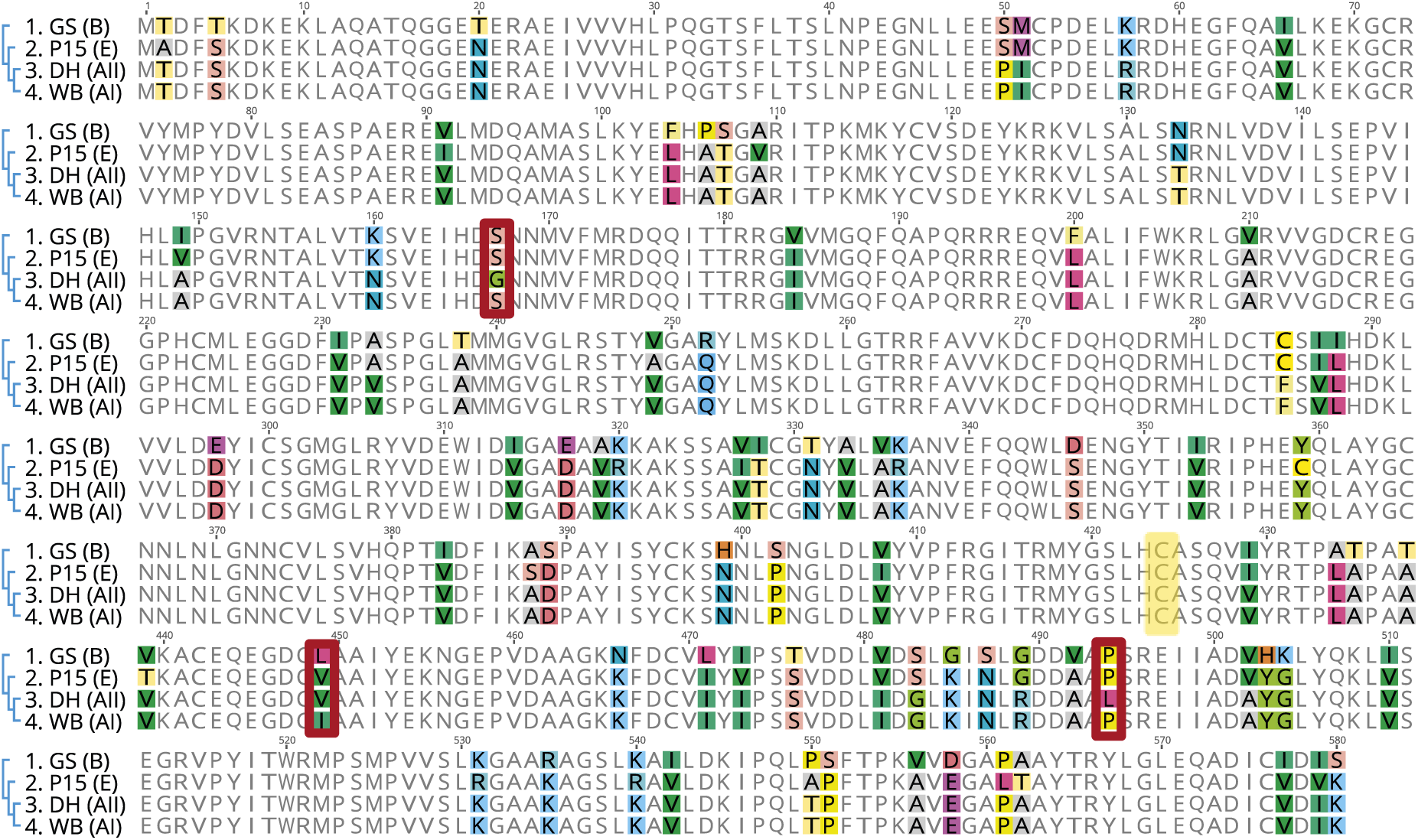
Variation in the amino acid sequences of ADI from indicated *G. duodenalis* reference isolates. Variant amino acids of ADI from assemblages AI (isolate WB), AII (isolate DH), B (isolate GS), and E (isolate P15) were color-coded with random color selection to highlight the amino acid sequence variation. The cysteine at the active center at position 424 (boxed yellow) and the three differing amino acids (boxed red) for assemblage AI (WB) and AII (DH) sequences at positions 167, 449, and 494 are marked.

**Figure 3.**
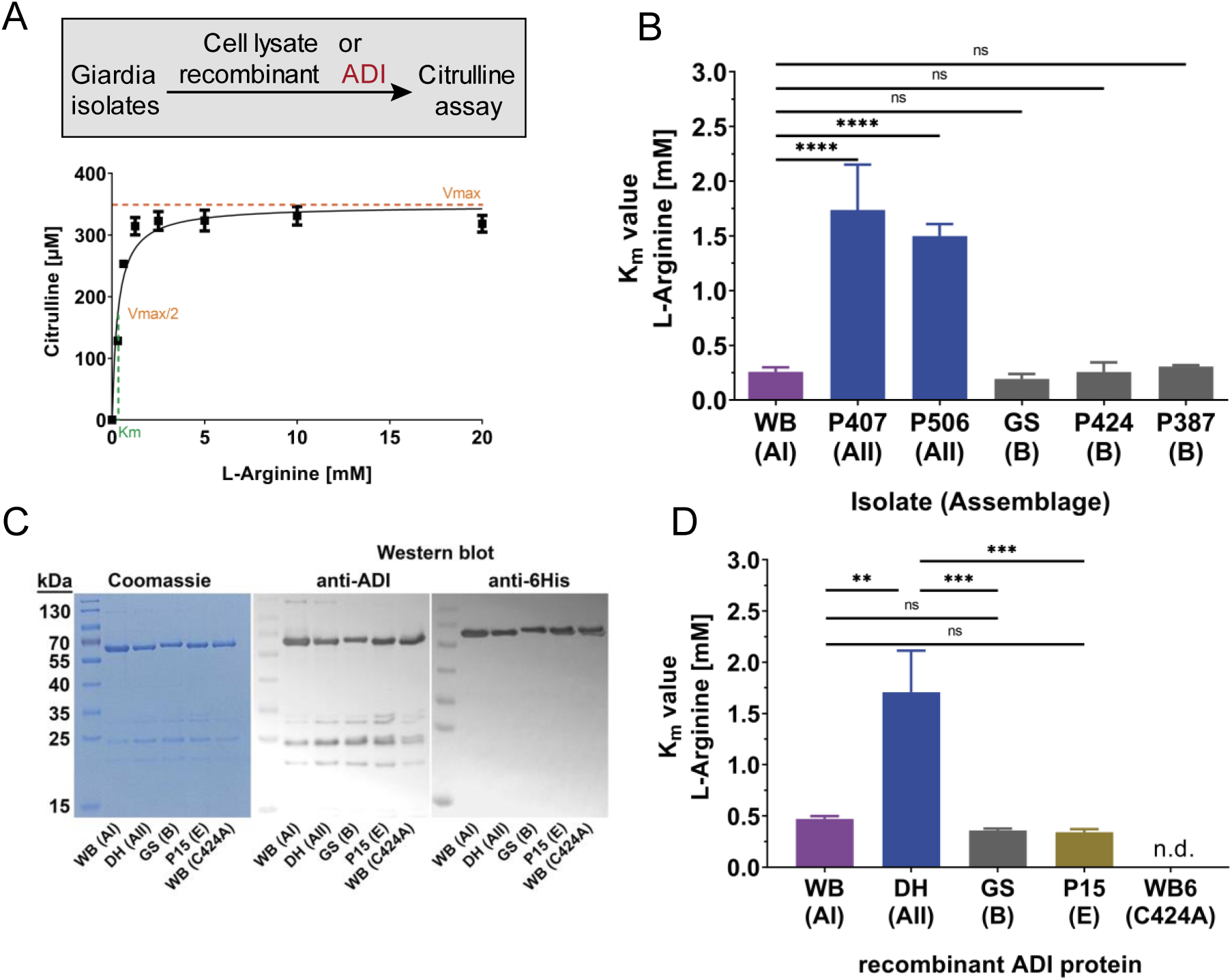
Lower substrate affinity of ADI from the assemblage AII isolates as compared to assemblages AI or B isolates. (A) Schematic representation and experimental example for the determination of the ADI *K*_m_ value for arginine. For determination of Michaelis-Menten kinetics for the respective *G. duodenalis* assemblage type ADI proteins, either the cell lysate or the recombinant ADI proteins were incubated with increasing concentration of the arginine substrate and formation of citrulline was quantified. (B) *K*_m_ values (mean ± SD) of ADI activity in lysates of *G. duodenalis* trophozoites demonstrate the significantly lower substrate affinity of ADI_AII_ enzymes from the assemblage AII isolates as compared to ADI_AI/B_ from assemblage AI and B isolates. Statistical testing was done using one-way ANOVA and Tukey post hoc tests on data of three independent experiments, each performed in triplicates (****, p < 0.0001; ns, not significant). (C) Recombinantly expressed 6His-tagged ADI proteins from assemblages AI (isolate WB), AII (isolate DH), GS (isolate GS), E (isolate P15) and an inactive ADI_AI_ (C424A) were affinity purified and analyzed by SDS-PAGE with Coomassie Blue staining as well as by Western blot analysis using a polyclonal alpaca anti-ADI serum [20] or an anti-6His antibody. (D) *K*_m_ values for each of the recombinant ADI proteins are shown. Protein sequences of ADI enzymes in lysates of assemblage AII isolates P407 and P506 in (C) are identical to the sequence of recombinant enzyme deduced from reference isolate DH. *K*_m_ values are presented as the mean ± SD of three independent experiments in triplicate. Statistical analyses shown used a one-way ANOVA and Tukey post hoc test, (**, p < 0.01; ***, p < 0.001; ns, not significant).

In Figure 2, the protein alignment of ADI sequences predicted from reference strains that we retrieved from the public database giardiadb.org [26] is shown. As expected, over the complete ADI gene sequence length of 1743 bp (580 amino acids) the sequence of assemblage AII (DH isolate) only displayed three different amino acids as compared with assemblage AI (WB isolate) but differed at 34 and at 64 amino acid positions in comparison with assemblage E (P15 isolate) and B (GS isolate), respectively (Figure 2).

Strikingly, at the positions of the three amino acids that differed between assemblage AI and AII (S167G, I449V, P494L), the assemblage B and E sequences are identical at two positions (S167, P494) compared to the sequence of assemblage AI. At position 449, assemblage B and E possess leucine and valine residues, respectively, with comparable biochemical characteristics to the cognate isoleucine at position 449 in the assemblage AI sequence (Figure 2).

To determine potential functional differences of ADI variants from different assemblages, we first generated cell lysates of *in vitro* cultured trophozoites from isolates of each of the different *G. duodenalis* assemblages and determined the substrate affinity of respective ADIs. By quantifying the product citrulline (using a standard assay), we determined the *K*_m_ values assuming Michaelis-Menten kinetics (Figure 3A). Overall, the *K*_m_ values of ADI_AII_ were approximately 5-fold higher than the *K*_m_ values from either the ADI_AI_ or ADI_B,_ which showed similar *K*_m_ values (Figure 3B).

### Recombinant ADI variants confirm substrate affinity differences between assemblages

The different substrate affinity of ADI_AI_ and ADI_AII_ was unexpected, as the two sub-assemblages share an overall high ADI sequence homology in comparison to ADI_B_ (see Figure 2). Notably, also based on whole genome level, assemblages AI and AII share sequence homology of approximately 99% [25, 28], whereas assemblages A and B only share approximately 70% sequence homology [33].

We therefore next asked whether the differences in enzymatic activity can be assigned to the three different amino acids in the underlying ADI_AI/AII_ sequences.

To confirm that the apparent *K*_m_ values were indeed due to the ADI enzymes and not due to unknown confounding activities in the parasite lysates, we recombinantly expressed ADI_I_, ADI_II_ and ADI_B_ in bacteria and determined the *K*_m_ values by using the different affinity-purified enzymes (Figure 3C, D). For further comparison, we also included a recombinant ADI from assemblage E (P15 isolate), an assemblage type that has been occasionally described in human infections, as well as an engineered inactive recombinant variant of ADI of WB isolate previously described [24, 34]. Overall, the *K*_m_ values of the recombinant proteins closely matched the *K*_m_ values observed in the trophozoite lysates from the respective assemblages. Again, the recombinant ADI_AII_ had a significantly decreased substrate affinity (Figure 3D). Of note, the ADI_E_ showed a similar *K*_m_ value as the ADI_AI_ and ADI_B_. The inactivate ADI protein expectedly showed no detectable substrate conversion (Figure 3D).

### Identification of key amino acid residues required for ADI substrate affinity

To test whether assemblage AII isolates in general carry the identified three amino acid differences compared to other assemblages (Figure 2), we amplified and sequenced the ADI genes from 14 different assemblage AII isolates established previously as cultures from stool samples of individual giardiasis patients [28, 32, 35]. All ADI_AII_ sequences displayed the same three amino acid differences as the DH isolate as compared to ADI_AI_ (Supplementary Figure S1). Most of the ADI_AII_ sequences (n=11) of the isolates studied were identical to the reference sequence of the DH isolate, but three patient isolates had one additional SNP at a different site in the gene sequences (Supplementary Figure S1).

In a similar manner, we sequenced the ADI_B_ genes derived from cultures of assemblage B patient isolates to confirm the conservation of the three specific ADI variant amino acid residues. *G. duodenalis* parasites are diploid organisms and possess a tetraploid genome, which leads to different degree of allelic sequence heterozygosity. Because assemblage B isolates often possess a high degree of ASH leading to ambiguous sequences using Sanger sequencing, we first cloned the ADI sequences from PCR products into a standard cloning vector and then analyzed 10 clones for each patient isolate. Altogether, for the 15 isolates tested, we identified 17 different alleles encoding the ADI_B_ protein (Supplementary Figure S2). In all ADI_B_ alleles, the positions of the three amino acids S167, L449 and P494 were conserved and identical to the GS reference isolate. Thus, our extensive analysis of ADI sequence variation in the various assemblages supports the existence of two specific amino acid mutations in the ADI_AII_ protein (at positions G167, L494) that are likely responsible for the functional differences in *K*_m_ (Figure 3B, 3D; Supplementary Figure S1, S2, S3).

To assess this hypothesis, we consecutively mutagenized the ADI encoding sequence of assemblage AI (WB isolate) at the three differing amino acid residues (ADI-167, ADI-449, and ADI-494) to match the assemblage AII ADI amino acid sequence. The eight modified ADI protein variants were recombinantly expressed and the *K*_m_ value determined for each variant. Of the eight possible variants, the two positions 167 and 494 (ADI_AI_ S167G and P494L) were sufficient to cause the functional differences in substrate affinity observed between ADI_AI_ and ADI_AII_ proteins (Figure 4A). Single changes in any of the three positions had no or only marginal effects on substrate affinity. Also, the amino acid exchange I449V had no significant effect on the substrate affinity of the ADI enzyme.

**Figure 4.**
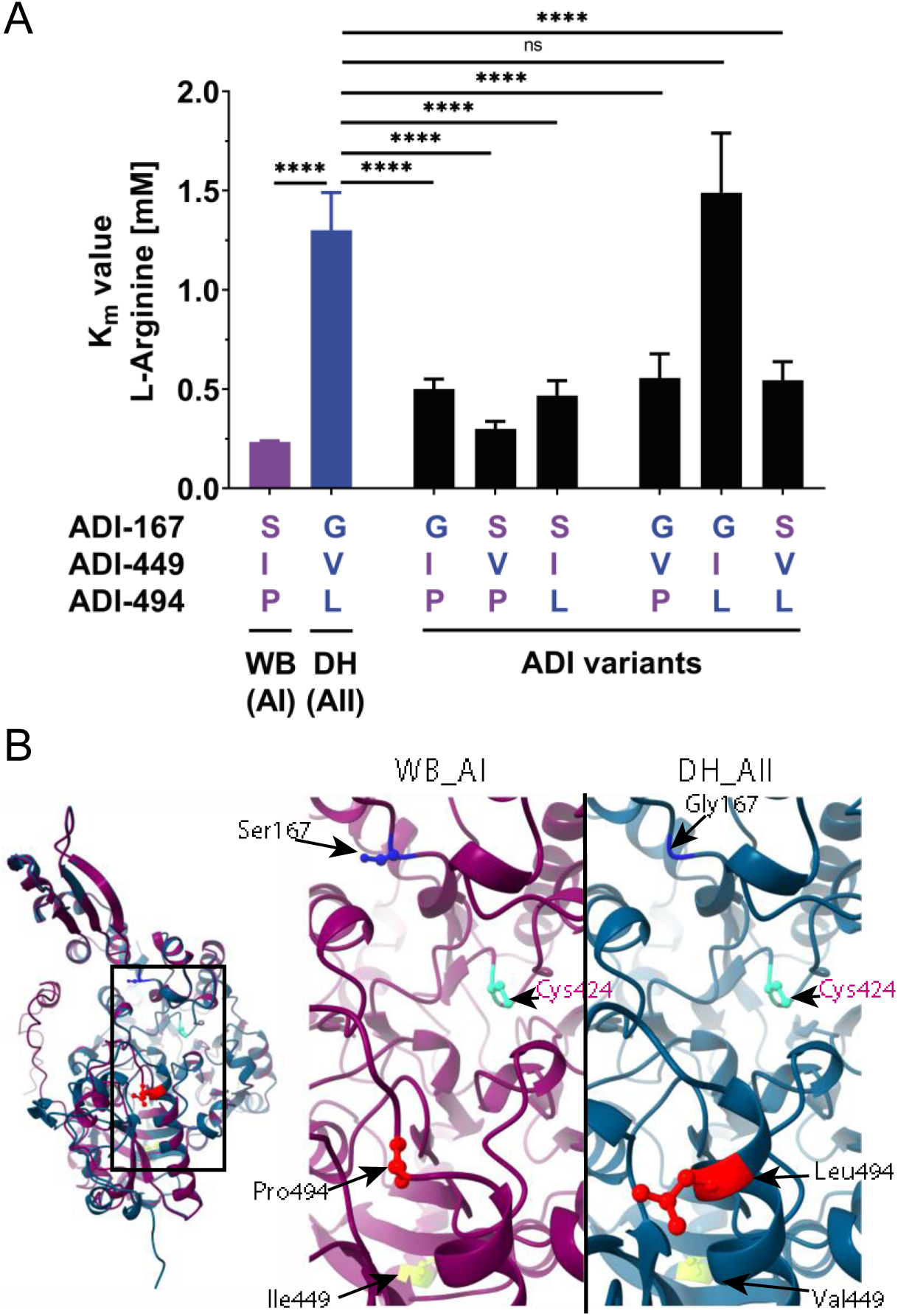
Two distinct amino acid residues near the active center confer the lower substrate affinity of ADI from the assemblage AII as compared to assemblage AI. (A) The arginine *K*_m_ values for recombinant ADI AI variants at amino acid position 167, 449, and 494 representing either AI or AII sequence residues were compared to unmodified assemblage AI (WB) and AII (DH) isolate ADI proteins. Mean ± SDs of 3 independent experiments with triplicates for each protein version are shown and analyzed by 1way ANOVA and Tukey post hoc to test for significance against *K*_m_ value of AII-type ADI (****, p < 0.0001; ns, not significant). (B) A structural comparison is presented for AlphaFold models of ADI_AI_ (isolate WB, pink) and ADI_AII_ (isolate DH, blue). The overlay shows the overall structural similarity of both ADI variants (left). The right panel highlights the comparative detail of the variant amino acid differences with the cysteine at the active center (pink).

Next, we retrieved the predicted protein structure for ADI of the reference strains WB (AI) and DH (AII) from AlphaFold 2 [36] and Chai-1 [37] and compared the structures using ChimeraX to reveal the possible structural implications of the amino acid positions ADI-167 and ADI-494 (Figure 4B, Supplementary Figure S4). Expectedly, ADI structures from assemblage AI and AII aligned well, particularly around the active center. This structural analysis, however, might predict that both variants, ADI-G167 and ADI-L494 as present in AII, likely may have a structural impact nearby the active center, whereas position 449 does not (Figure 4B). Further structural analysis with arginine positioned in the active center and only considering the mutations at position 167 and 494, highlight the position of the two mutations in loop structures (Supplementary Figure S4).

### An essential role for arginine metabolism in encystation in G. duodenalis assemblages

In the gut, completion of the *G. duodenalis* life cycle relies on successful differentiation of the trophozoite to the environmentally resistant cyst via a highly regulated developmental process [20, 23, 38, 39]. This encystation is a highly energy-dependent process and includes incomplete division, specific assembly of the endomembrane system and the cyst wall. Previous studies have suggested a regulatory role of ADI and the ADH pathway for ATP production required for encystation [15].

We therefore hypothesized that the lower ADI substrate affinity of ADI_AII_ may also affect the overall encystation efficiency of AII isolates. Thus, using standard encystation protocols, we compared the encystation of isolates WB (AI) and P407 (AII). The encystation efficiency of AI trophozoites (∼55%) was indeed significantly higher than that of AII trophozoites (∼30%) (Figure 5A).

**Figure 5.**
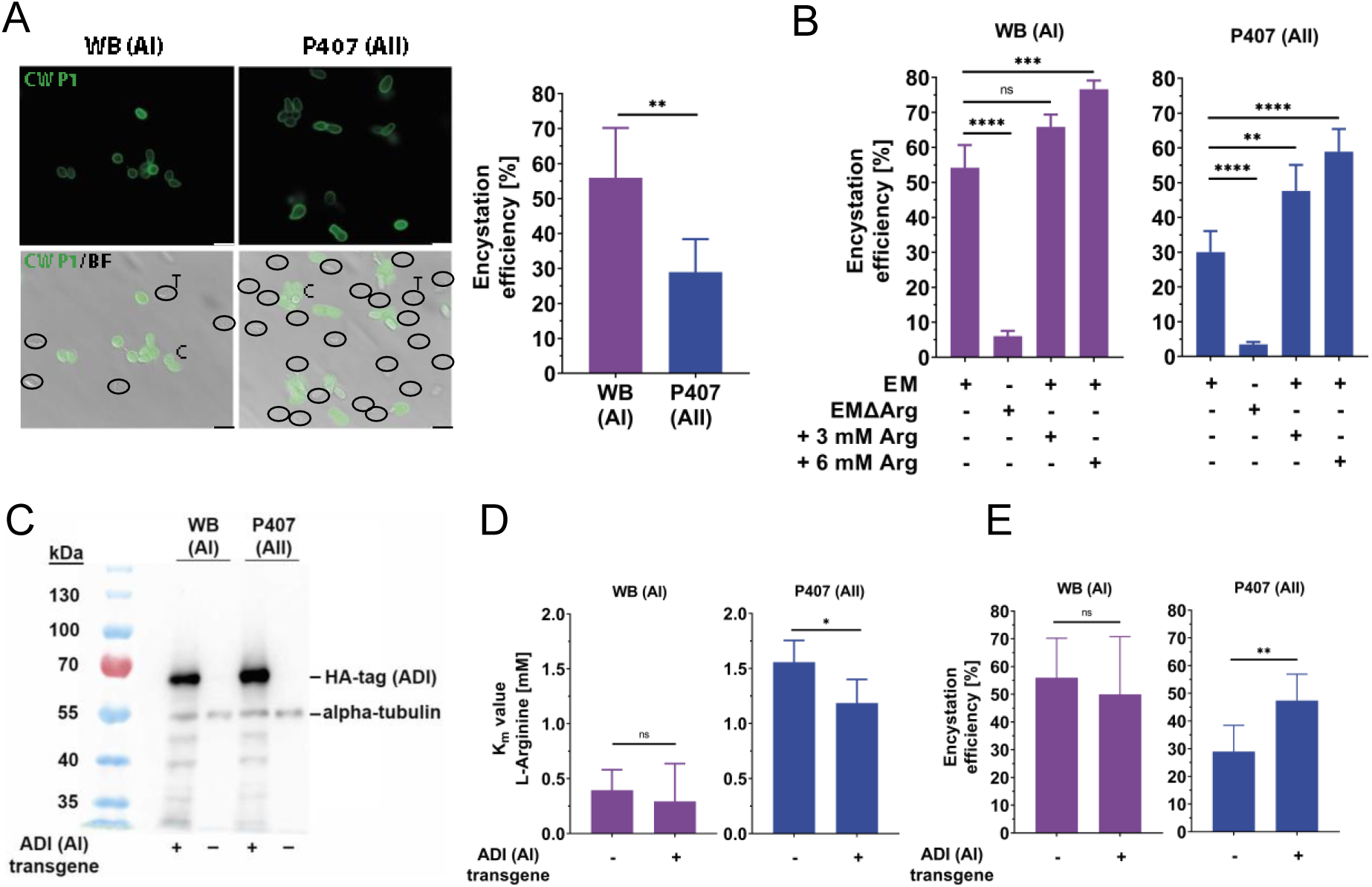
Encystation efficiency depends on arginine availability and ADI affinity. (A) Differences in the efficiency of in vitro encystation compared between the assemblage AI isolate WB and assemblage AII isolate P407. Representative bright field (BF) and immunofluorescence images using anti CWP1-antibody (green, left) illustrate the lower encystation efficiency of P407 in comparison to WB isolate (graph, right). CWP1-negative trophozoites are circled for better visualization. Scale bar = 10 µm. Encystation efficiency values compared between isolates show the mean ± SD of three independent experiments in triplicates (statistical analysis by unpaired t-test, **, p < 0.01). (B) The in vitro encystation efficiency in both assemblage types AI and AII is drastically reduced in arginine-depleted encystation media (EMΔArg) and efficiency is increased by supplemented arginine. The arginine concentration in EM was calculated to be approximately 3 mM. Values plotted represent the mean ± SD of three independent experiments in triplicates, and statistical analyses were performed using a one-way ANOVA and Tukey post hoc test (**, p < 0.01; ***, p < 0.001; ****, p < 0.0001; ns, not significant). (C) Western blot analysis was performed using an anti-HA antibody to detect recombinant HA-tagged AI-type ADI expression constructs transfected into both P407 (assemblage AII) and WB (assemblage AI) isolates. A pan alpha-tubulin antibody was used as a loading control. (D) The relative arginine *K*_m_ of ADI activity in lysates is shown for both transgenic WB (AI) and P407 (AII) trophozoites. Mean ± SD of three independent experiments in triplicates are indicated and statistical analysis performed as in B (*, p < 0.05). (E) Increased encystation efficiency in transgenic P407 (AII) expressing additional AI-type ADI as compared to WB (AI) control isolates. Mean ± SD encystation efficiencies for three independent experiments in triplicate are shown with statistical testing as performed in B (**, p < 0.01).

To confirm the essential role of arginine metabolism in encystation, we first enzymatically depleted arginine from the medium using the recombinantly expressed ADI_AI_ protein and then compared the encystation of AI vs. AII isolates (Figure 5B). Encystation efficiency was dramatically decreased for both AI WB and AII P407 isolates in depleted medium, with AII P407 cells showing an residual encystation efficiency of about 5% overall (Figure 5B). In contrast, supplementation of an additional 3 or 6 mM arginine to standard encystation medium (modified TYI-S-33, [40]), which already contains approximately 3 mM arginine, led to an increase in encystation to 75% for AI WB and 60% for AII P407 (Figure 5B).

To further investigate the impact of the ADI_AI_ and ADI_AII_ enzyme substrate affinity in the two variants and the link of arginine metabolism to the encystation, we generated transgenic WB and P407 parasites expressing an integrated HA-tagged, type ADI_AI_ gene copy under the endogenous promoter using previously described methods [41]. Both strains had stably integrated the ADI_AI_ transgene into the genome and expressed similar amounts of the HA-tagged type ADI_AI_ protein (Figure 5C). The *K*_m_ values in lysates of both wild-type and transgenic WB parasites (assemblage AI) revealed similar *K*_m_ values (Figure 5D). In contrast, AII (P407) parasites expressing the type ADI_AI_ enzyme had a lower *K*_m_ value. This indicates that the expressed type ADI_AI_ transgene improved the resulting affinity of the ADI protein in the AII (P407) transgenic parasites, which also expressed their endogenous type ADI_AII_ protein (Figure 5D). In line with this result, the transgenic AII strain encysted significantly more efficiently compared to the wild-type P407 strain, whereas encystation of the transgenic AI strain was not altered (Figure 5E).

For assemblage B isolates, no *in vitro* encystation protocol is available. Thus, to test the generality of the arginine dependency for encystation in different assemblages in the host, we infected mice fed either a diet with arginine-containing or arginine-free food pellets with assemblage B GS/H7 parasites. While both groups did not differ regarding trophozoite loads in the duodenum at day 7 post-infection, mice fed the arginine-depleted diet shed significantly fewer cysts in their feces (Supplementary Figure S5). Thus, studies both *in vitro* and *in vivo* support that arginine availability as well as ADI enzyme activity are indeed key factors for *G. duodenalis* cyst formation.

### ADI gene disruption confirms the essential role of arginine metabolism for encystation

Finally, we used a new genetic strategy for *G. duodenalis* [42] to create an ADI knockout mutant to prove that a functional ADI enzyme is essential for efficient encystation in *G. duodenalis.* Using CRISPR/Cas9 we created an ADI knockout mutant (ADIKO) strain by consecutively disrupting all four ADI alleles in the reference strain WB using two antibiotic resistance markers [42]. The absence of ADI protein expression in ADIKO strains was confirmed in clones by both, Western blot (Figure 6A) and the lack of ADI activity in ADIKO cell lysates (Figure 6B). While two ADIKO clones showed sustainable growth in standard growth medium, each had decreased generation times compared to wild type parasites (∼10 hours for ADIKO versus ∼7 hours for wild-type). Strikingly, cyst production in ADIKO parasites in standard encystation medium showed a drastic reduction of cyst production in comparison to wild-type parasites.

**Figure 6.**
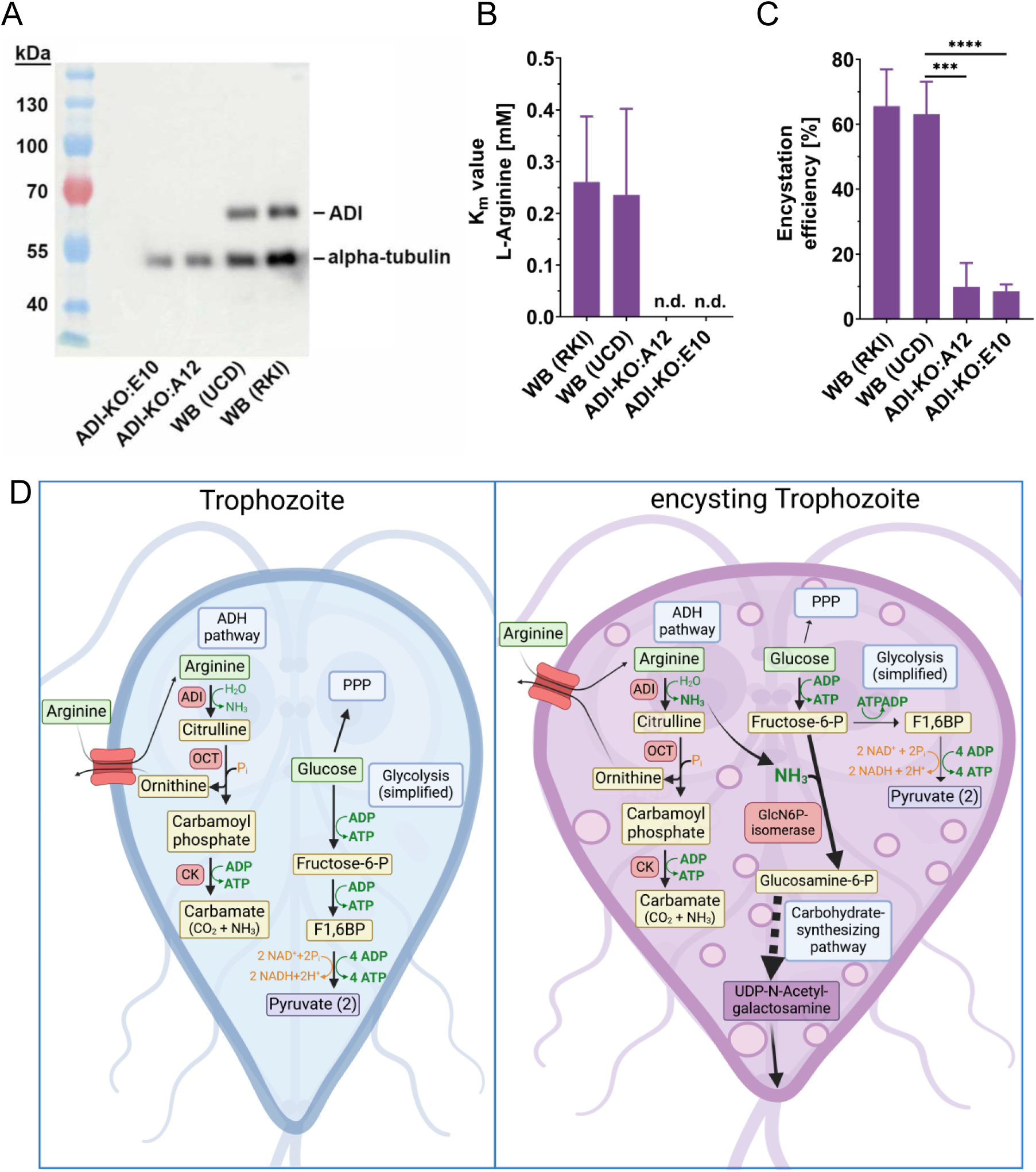
Arginine deaminase activity essential for efficient encystation in *G. duodenalis.* Quadruple allele ADI knockouts were generated using a CRISPR/Cas9 approach (see Methods and [39, 61]). (A) The lack of ADI expression from two independent ADI knock-out clones (E10, A12) in the assemblage AI (WB) isolate was verified by Western blot analysis using a polyclonal anti-ADI antibody and a pan alpha-tubulin antibody as loading control. The parental WB (UCD) strain as well as another WB strain (RKI) maintained in a different laboratory were included as ADI-expressing controls. (B) ADI enzymatic activity was not detectable (n.d.) in lysates of both ADI-KO clones while both WB control strains (“RKI” and “UCD”) showed similar *K*_m_ values. Mean ± SD *K*_m_ values from three independent experiments in triplicates are given. (C) The encystation efficiency (see Figure 5A) was drastically reduced in both ADI-KO clones (mean ± SD of three independent triplicate experiments). Statistical tests were performed by using one-way ANOVA and Tukey post hoc test (***, p < 0.001; ****, p < 0.0001). (D) Cartoon of selected, simplified biochemical pathways for arginine and glucose in trophozoites and encysting trophozoites. In the trophozoite stage, the ADH pathway and glycolysis both represent important, redundant pathways for ATP production in this rapid growing stage. Glucose is also an important reactant for alternative pathways, such as the pentose phosphate pathway (PPP), to generate pentoses. In the encysting trophozoite, glucose is required for generation of the major carbohydrate cyst wall component UDP-N-acetylgalactosamine via a carbohydrate synthesizing pathway [43]. Other glucose-dependent pathways, such as glycolysis or PPP, are therefore downregulated. In this scenario, the ADH pathway is fundamental for energy (ATP) production. In addition, glucosamine 6-phosphate isomerase (GlcN6P-isomerase), which is the first enzyme in the carbohydrate synthesizing pathway, requires ammonium (NH_3_) to generate glucosamine-6-phosphate from fructose-6-phosphate.

## Discussion

*G. duodenalis* follows a simple life cycle, in which phases of massively proliferating trophozoites populating the small intestine proceed to differentiation into cysts, the transmission stage excreted with feces. Virulence, i.e., the capability of this purely luminal gastrointestinal pathogen to cause disease, is linked to parasite numbers. This was shown by the infection dose-dependent breakdown of barrier functions in our recently developed human duodenal organoid-derived model of infection [11]. Furthermore, epidemiological data link symptomatic disease to test positivity in less sensitive diagnostic methods [3].

Throughout the life cycle of these microaerophiles, energy in the form of ATP is derived from substrate level phosphorylation pathways. In *Giardia*, as in eukaryotes in general, glycolysis and pyruvate metabolism yield two ATP for every glucose metabolized. In addition, these parasites possess the ADH pathway [43, 44] that can generate one ATP for every catabolized arginine molecule (Figure 6D). Transcriptomic and proteomic studies suggest stage-dependent metabolic adaptations that include regulation of those energy-producing pathways [20–23]. Biochemical data support the concept that ATP generation by the ADH pathway is particularly relevant for life cycle phases during which funneling of glucose into anabolic pathways is required. For example, the growth promoting effect of arginine is important in phases of high biomass generation, i.e., fast trophozoite growth [43], when glucose may have to feed into the pentose phosphate shunt. Another example, as shown in the present work, is the effect of arginine concentration and ADI promoting cyst formation when glucose is required for generation of N-acetlygalactosamine-containing cyst wall components during encystation [45]. The key role of ADI as the first enzyme of the ADH pathway in the encystation process and the relevance of allelic forms as shown here (see also below) now firmly consolidates the proposition that ADI indeed is a virulence factor, i.e., a quantitative trait linked to pathogenicity and transmissibility. It is intriguing that the enzymes of the ADH pathway, i.e., ADI, OTC, and CK, were likely acquired by *Giardia* from bacteria by horizontal gene transfer [19]. Conceptually, this may be considered akin to the acquisition of pathogenicity islands in bacterial pathogens [46].

### The amino acid arginine is a key component of energy metabolism in Giardia

Arginine metabolism is a key component of parasite energy production during both life cycle stages, and it is also proposed to compete for arginine with host epithelium [15, 17, 18, 43]. ATP production by arginine metabolism is preferred over glycolysis in trophozoites, and arginine is depleted in *Giardia* growth medium during *in vitro* culture [43]. During encystation, arginine metabolism has been proposed to act as an alternative ATP source due to the requirement of metabolites from glycolytic intermediates, which are required for production of cyst wall components that may be diverted from glycolysis and glucose metabolism [45]. In support, genes of the ADH pathway are among the genes highly expressed in the trophozoite stage and this high expression level remains largely unchanged during encystation. Only in the later cyst stages, transcripts and protein abundances are lower [20–23].

Parasite arginine metabolism also limits or modulates host immune responses in giardiasis. In the anoxic gut, competition for arginine with cells of the host epithelium [16–18] or microbiome [47] may induce both host pathophysiology and immune modulation [10]. Thus, competition for arginine between the parasite and host during either trophozoite metabolism and/or encystation may account for host pathophysiology, but needs further investigation using appropriate host cell-pathogen systems, e.g., in our recently described model based on stem cell-derived intestinal organoids [11, 48].

### The ADI pathway has both definitive catabolic and anabolic metabolic functions during Giardia encystation

The ADI pathway plays an important catabolic role in parasite metabolism. Here, using both biochemical and new genetic approaches in *Giardia* research, we demonstrate that a functional ADI enzyme is essential for efficient encystation, thereby highlighting how this energy generating pathway is required for transmission of this ubiquitous pathogen. Both, genetic knock-out of all four ADI alleles or depletion of the ADI substrate arginine in the culture medium led to a drastic inhibition of cyst production. While neither one of these processes is lethal for the trophozoites, yet each one reduces the trophozoite growth rate. Thus, *Giardia* uses redundant mechanisms for energy production by substrate level phosphorylation in the trophozoite stage, but requires the ADH pathway to fuel the encystation process efficiently. Note, previous attempts to produce functional ADI-knock outs using siRNA knock-down technologies were not successful [34], indicating parasite metabolic adaptation is needed to outgrow the KO parasites.

In addition, products of the ADH pathway may also play an important role in anabolic metabolism during encystation. Encysting *Giardia* parasites use glycolytic intermediates for the production of UDP-N-acetylgalactosamine, which is an important precursor for cyst wall components [45]. Thus, encystation may deplete these metabolic intermediates and limit energy production by glycolysis during encystation. In support, a number of groups have produced valuable inventories of transcripts and proteins for studying encysting WB parasites [20–23]. The most comprehensive study to date [22] confirms in great detail a higher abundance of enzymes required to synthesize the N-acetylgalactosamine. These components remain at a high level in encysting cells and cysts while glycolytic proteins, in particular those associated with ATP production, are downregulated.

To synthesize UDP-N-acetylgalactosamine from endogenous glucose, *Giardia* needs to generate glucosamine-6 phosphate from fructose-6 phosphate using glucosamine-6 phosphate isomerase. This is a cytosolic enzyme with aminase activity; induced in encysting parasites and also dependent on NH_3_ as a further substrate [45] (see Figure 6D). Thus, we postulate that ADI not only provides the citrulline-intermediate for the ADH pathway-dependent ATP generation but also increases the ammonia/ammonium concentration within the encysting trophozoite. Thereby, anabolic and catabolic functions of the ADH pathway fuel glucosamine-6 phosphate synthesis and eventually cyst wall formation. ADI, OCT and CK are progressively less abundant in maturing cysts, as also confirmed for ADI by our immunofluorescence analysis (Figure 1B). Furthermore, the progressive deposition of the cell wall is thought to act as a diffusion barrier [49] and may limit arginine influx and thus act as a natural negative feedback loop to signal that cyst wall generation is completed. This could explain the dependency of encystation on the substrate affinity of ADI.

### Variant arginine substrate affinity of ADI may impact encystation efficiency and host metabolic interactions in different G. duodenalis assemblages

This work provides the first direct functional evidence for genetically determined virulence in *G. duodenalis*. Moreover, we show that arginine substrate affinity of the ADI enzyme varies between different *Giardia* assemblages. The sharp distinction in ADI substrate affinity between *G. duodenalis* assemblage AI and AII was unexpected, as ADI gene similarity reaches 99.6% with only three amino acid differences on protein level between the two groups. We identified the two conserved amino acid residues in the ADI_AII_ sequence that are responsible for the functional difference. Strikingly, ADI_B_, with only ∼89% protein sequence homology to ADI_AI/AII_ [1, 28, 30, 33, 50], provides comparable biochemical function as ADI_AI_. It possesses the identical two conserved amino acid residues in question as type ADI_AI_. The structure prediction showed the positioning of the two functional mutations nearby the active center, but not in direct vicinity. In a recent study Li et al, by combinatorial mutating ADI from Enterobacter faecalis, showed similar modulation of catalytic activity in mutations in the loop structures nearby active center [51]. Further structural studies in combination with functional assays comparing ADI from different organisms in future will be needed to unravel underlying structural details responsible for the substrate affinity.

Earlier studies showed the presence of an extremely efficient arginine/ornithine antiporter on the plasma membrane of *G. duodenalis* with *K*_m_ values as low as 15 µM [52]. Until now, there is no evidence for efficient arginine biosynthesis in *G. duodenalis* and efficient uptake seems essential. For comparison, the most efficient arginine transporter (CAT-1) in human cells has a *K*_m_ value of 100-160 µM [53] -- approximately 10-fold less efficient. This difference in arginine transporters between parasite and host argues for a strong competition for free arginine [15–18]. Importantly and in contrast to e.g. *Trichomonas* parasites, there is no intracellular free arginine pool detectable in *G. duodenalis*, which likely is a consequence of the highly active ADH pathway [54].

Considering these observations, different *G. duodenalis* assemblages have different arginine affinities, with likely consequences for parasite metabolic variation during encystation and host-interactions. Specifically, the *K*_m_ values of ADI of the assemblage AI or B type enzyme ensure more optimal arginine consumption under conditions, where lower arginine amounts are available, while the assemblage AII type enzyme likely performs sub-optimally under this assumption. Furthermore, we not only show the importance of ADI allelic types for encystation, but also have identified the two SNPs and deduced amino acid sequences in the ADI loci of human specific assemblage AII isolates that define this functional difference in arginine substrate affinity between ADI proteins in different assemblages.

Recent molecular epidemiological studies indicate that assemblage AII represents an anthroponotic assemblage type only found in humans, whereas assemblage AI and B represent potentially zoonotic assemblage types populating humans and a wide range of animals [1, 7, 8]. Our analysis of allelic ADI variants of patient isolates indicates that the two amino acid residues distinguishing ADI of anthroponotic from zoonotic assemblages are evolutionary conserved within the assemblage groups.

Differences in sequence and function of ADI_AII_ could therefore be a consequence of host adaptation. Host dietary requirements for arginine are highly species-specific, and while such requirements are well-studied in livestock species and companion animals, dietary requirements for arginine are not well understood for humans [55]. With respect to parasite ADI evolution, selection pressure in different hosts may act directly on parasite fitness with respect to the completion of the transmission stage (e.g., encystation). Hence, differences in parasite metabolism amongst the different *G. duodenalis* assemblages, such as the functional differences in ADI arginine affinity, could contribute to the wider zoonotic distribution of some *G. duodenalis* assemblages (AI and B) as compared to the narrow anthroponotic distribution of the AII assemblage.

## Material and Methods

### Trophozoite culture

Trophozoites of the *G. duodenalis* strains WB (WB-C6, ATCC 50803) and GS (GS-H7, ATCC 50581) were propagated in TYI-S-33 medium, supplemented with adult bovine serum, in 11 ml slanted screw capped culture tubes as previously described [40]. For passaging, confluent culture tubes were incubated on ice for 20-30 minutes to detach trophozoites, and a sufficient proportion of trophozoites was transferred into new tubes filled with fresh medium. *G. duodenalis* parasites from patients were isolated from stool samples by *in vitro* excystation following the protocol of Rice and Schaefer [56] and cultured in TYI-S-33 medium as previously described [28, 32, 35]. Briefly, cysts were enriched from stool samples by sucrose gradient flotation, excysted in vitro and cultures established through limiting dilution in 96-well plates under anaerobic conditions. Growing isolates were transferred into culture tubes for routine culture.

### Parasite encystation

Encystation was induced according to previously published protocols using modified TYI-S-33 medium, containing higher bile concentrations and an increased pH value [57, 58]. In brief, trophozoites were grown to logarithmic phase between 60-80% confluency in culture tubes or 12-well plates with normal TYI-S-33 medium containing 0.5 mg/ml bovine bile (Sigma, B8381) at a pH of 7.0. For encystation, normal growth medium was replaced by the same amount of warm encystation medium, i.e. TYI-S-33 medium containing 10 mg/ml bovine bile and adjusted to a pH value of 7.85, and incubated for 24 hours at 37°C, unless stated otherwise. Trophozoites that were cultured in cell culture plates were put under oxygen-restricted growth conditions using Anaerogen Oxoid jars containing appropriate reaction bags (Thermo Fisher, AnaeroGen #AN0025).

To quantify encystation, cultures were put on ice for 20 min to detach all parasites (trophozoites, encysting trophozoites and cysts) from the cell culture plastic. Parasites were collected, pelleted at 900 *g*, 4°C and fixed with 4% paraformaldehyde in PBS for 15 minutes at room temperature. After fixation, cells were washed twice and finally resuspended with PBS. Subsequently, cells were treated 30 minutes at room temperature with 1x Giardi-a-glo antibody labeled with FITC (Waterborne Inc.) to detect cysts and nucleic acid stain propidium iodide (final concentration of 5 µg/ml) for counterstaining. In some experiments, cells were counterstained with Troph-o-glo antibody labeled with Cy3 (Waterborne Inc.) to visualize trophozoites and Hoechst 33342 stain at a final concentration of 2 µg/ml to stain nuclei. Finally, cells were washed with PBS and counted using counting chambers under conventional fluorescent microscope (Zeiss Axioscope). Efficiency of encystation was calculated by dividing number of cysts with total cell count (cysts and trophozoite stages).

### Animal infection

Standard *in vitro* encystation protocols are not efficient for assemblage B parasites. We therefore used an established infection protocol in mice [59] to determine the role of arginine on encystation in vivo. In brief, 6-10 week-old C57/BL6 mice (female and male) were randomly divided into experimental groups and pretreated for seven days with antibiotics in drinking water ad libitum, containing 1.4 g/L Neomycin (Biomol, Germany), 1 g/L Ampicillin (Merck, Germany) and 1 g/L Vancomycin (Merck, Germany). At the same time, the diets of the experimental groups were exchanged towards “arginine-free” and “control diet” (EF Crystalline AA Arginine free and EF Crystalline AA control diet, ssniff GmbH, Arnsberg, Germany). Animals were orally infected by gavage with 5 x 10^6^ trophozoites derived from an in vitro culture of the *G. duodenalis* GS clone H7 (ATCC 50581, assemblage B). Feces were collected daily from day 4-7 post infection and cyst shedding was determined in suspended feces (100 mg feces per ml water) by immunofluorescent microscopy using Giardi-a-glo antibody (Waterborne inc). On day 7 post infection, animals were sacrificed and a 3 cm piece of the small intestine, starting at 2 cm from the pylorus, was cut out and frozen in 1 ml RNALater solution (Qiagen, Germany) at -70° C until further processing. For DNA extraction, intestinal tissue was first homogenized in a Pecellys 24 tissue homogenizer (Bertin Instruments, France) using a bead-mixture (50/50% 1 mm and 0.1 mm sharp-edged silicon carbide beads from Bio Spec Products Inc, pre-treated at 180°C for 2.5 h) and RLT buffer from Qiagen RNeasy Mini kit (Qiagen, Germany) with 1% RNase free 2-mercaptoethanol (Merck, Germany). Homogenates were centrifuged for 3 minutes at 21.500 g and supernatants were used to isolate DNA with the NucleoSpin Tissue kit (Macherey-Nagel, Germany) according to the manufacturers’ recommendations. Quantification of *G. duodenalis* parasites was done by a published Taqman qPCR protocol targeting the rRNA gene from *G. duodenalis* [60]. An internal amplification control was included to exclude potential PCR inhibition [61]. Genome equivalents were calculated from a standard curve produced from DNA of a known number of in vitro grown GS/H7 isolate. The work has been approved by local authorities (Landesamt fuer Gesundheit und Soziales, Berlin, Germany, license number G_0277-17).

### Localization of ADI by immunofluorescence analysis

Localization of ADI protein was analyzed in trophozoites, encysting trophozoites and cysts of WB isolate. Cysts were separated from encysting trophozoites 24 hours after induction of encystation by centrifugation at 650 *g* for 10 minutes at room temperature. Cysts were collected and washed once with H_2_O and stored in H_2_O at 4°C for 48 hours and then fixed with 4% paraformaldehyde for 15 minutes at room temperature and washed twice with PBS. Remaining encysting trophozoites were detached by putting the tube on ice for 20-30 minutes and then cells were pelleted for 10 minutes at 900 *g* at 4°C. Trophozoites were harvested from a routine culture in logarithmic phase as stated above. Trophozoites and encysting trophozoites were washed two times with PBS and then fixed with 4% paraformaldehyde for 15 min at room temperature. After further two washing steps with PBS, cells were stored at 4°C until usage. For immunofluorescence staining, cells were either immobilized on poly-L-lysin pretreated chamber slides (ibidi; Germany) or stained in suspension. First, cells were permeabilized with 0.25% TritonX-100 in PBS containing 0.75% glycine for 30 minutes and then blocked with 2% BSA in PBS for further 2 hours. Primary antibodies were diluted in blocking buffer and samples were incubated overnight at 4°C. Primary antibodies used: alpaca antiserum generated against recombinant ADI from WB isolate [24] in a dilution of 1:100. After 3 washing steps with blocking buffer for 10 minutes each, secondary antibodies (goat anti lama DyLight650, 1:300; Bethyl laboratories) and directly labeled antibody Giardi-a-Glo (FITC) were diluted in blocking buffer and incubated for 2 hours at room temperature. Cells were washed again 3 times for 10 minutes each. Finally, DNA was stained with Hoechst 33342 at a final concentration of 2 µg/ml for 30 minutes and cells were imaged in mounting medium (ibidi) at a Leica Mica microscope (Leica).

### Sequencing of isolate-specific G. duodenalis ADI alleles

The *G. duodenalis* ADI coding sequences of 15 assemblage AII isolates were amplified from genomic DNA by PCR using specific primers (5’-CTGACAAGCACTTCATTTACTG-3’ and 5’-CGGCGGGGGCCGGTGCTTTG-3’) and the Q5 PCR kit (New England Biolabs). PCR fragments were purified with the DNA Clean and Concentrator^TM^ kit (ZymoResearch). Sequencing was performed using the same amplification primers and two additional ones (5’- GTCCGCAACACGGCTCTCGTTAC-3’ and 5’-CCGAGGCGCTTCCAGAAGAT-3’) to retrieve the complete ADI coding sequence.

To determine the alleles of the ADI coding sequence of 15 selected assemblage B isolates, modified primers were used to amplify their ADI sequence (5’- CACAGCGTTTAATTTACATCTTATAAG-3’ and 5’CTAGACATAAACATCTCAATTATTTG-3’). The majority of assemblage B isolates are heterozygotes and we therefore cloned the PCR products into a standard cloning vector (CloneJET; ThermoScientific) and analyzed plasmids of 10 single bacterial clones per isolate. Plasmids carrying the ADI coding sequence were sequenced as described above.

Sequence files of both strands amplified by respective forward and reverse primers were finally analyzed with tools implemented in Geneious Prime software (Biomatters Ltd.) and CLC genomic workbench (Qiagen).

### Generation of recombinant proteins

The recombinant *G. duodenalis* ADI proteins corresponding to assemblage AI type GL50803_112103 and its enzymatically inactive mutant form were produced as described [24]. Assemblage AII, B as well as E encoding ORFs were designed based on their genomic sequence (Gene IDs DHA2_112103, GL50581_1575 and GLP15_4932; [26]) but codon optimized for expression in *E. coli* and custom ordered (GeneArt^®^, Invitrogen). The DNA fragments were cloned into the expression vector pASG-IBA35 (StarGate cloning, IBA GmbH) according to the manufacturer’s manual to produce a recombinant protein with an N-terminal His_6_-tag. Site directed mutagenesis using Q5 polymerase (New England Biolabs) was done according to the guidelines by the manufacturer, creating ADI with modified coding sequences, thereby transforming assemblage ADI_AI_ consecutively into assemblage ADI_AII_ in the pASG35 expression vector. The following amino acid positions and combinations thereof were modified (S167G, I449V, P494L, see Figure 2; 4A). After transformation of pASG-IBA35_ADI into *E. coli* DH5αZ1 cells, the expressed His_6_-tagged ADIs were purified by affinity chromatography followed by imidazole removal and concentration in PBS as described previously [24]. Protein concentrations were determined with a BCA protein assay kit (Thermo Scientific Fisher). Purified proteins were stored in aliquots at -70°C. Purity of recombinant proteins was controlled by SDS-PAGE and Coomassie staining. For Western blot analysis, the following antibody combinations were used: a commercial monoclonal mouse anti-6His antibody (1:2000, Qiagen) together with a peroxidase conjugated goat anti-mouse antibody (1:5000, Bethyl laboratories); an alpaca antiserum from an animal immunized with recombinant assemblage ADI_AI_ (1:1000) together with a peroxidase conjugated goat anti-lama IgG (1:10000, Bethyl laboratories). For detection, the Pierce ECL Plus substrate kit (ThermoFisher Scientific) was used and signals were recorded on a conventional documentation system (Vilbert Fusion FX6).

### Determination of enzymatic activity and enzyme K_m_ value

ADI activity and *K*_m_ values of recombinant, purified *G. duodenalis* ADIs were determined by colorimetric determination of citrulline formation as described [24, 52]. To determine the *K*_m_ value of native *G. duodenalis* ADI, lysates of trophozoites were prepared. Per strain, three cultivation flasks of confluent trophozoites were washed with PBS at room temperature and then placed for 30 min on ice. Cultures were harvested, combined and centrifuged (900 x *g*, 5 min, 4°C). After additional washing with ice-cold PBS, cell pellets were resuspended in 500 µl ice-cold PBS, trophozoites were counted in a Neubauer counting chamber and frozen at -80 °C. Subsequently, frozen trophozoites were thawed, exposed to three freeze/thaw-cycles using consecutive steps using liquid nitrogen and a 37°C water bath. Cell debris was removed by centrifugation for 20 min at 12000 x *g* at 4°C. Supernatants were collected and 10 µl lysate were directly or in appropriate dilution with 40 mM Hepes buffer (pH 7.0) used for the enzyme activity assay. To determine the *K*_m_ value, the substrate concentration was varied: 10 µl of different arginine concentrations, prepared by diluting 100 mM arginine stock solution with 40 mM Hepes buffer (pH 7.0), were applied to obtain final assay concentrations ranging from 0-20 mM. As negative control, 10 µl lysate was replaced by 40 mM Hepes buffer (pH 7.0) for each arginine concentration. *K*_m_ values were computed with GraphPad Prism 9 (GraphPAD Software, Inc.).

### Generation of transgenic and knock-out strains

For generation of assemblage AI and AII trophozoites carrying an additional genome integrated ADI_AI_ transgene, the ADI_AI_ gene (GL50803_112103), including its endogenous promotor, was PCR amplified from genomic DNA of *G. duodenalis* WB isolate using primer pairs (forward 5’GGGCCTAGGATGACTGACTTCTCCAAGGATA3’; reverse 5’ GGGGCATGCCTTGATATCGACGCAGATGTC3’) and Q5 PCR amplification kit (New England Biolabs). The PCR product was inserted by Gibson cloning procedures in vector pPAC-ORF16653-HA [62, 63] replacing ORF16653 with the ADI_AI_ gene, resulting in vector pPAC-ADI_AI_-HA. The vector was linearized using SwaI restriction enzyme and trophozoites of isolates WB (assemblage AI) and P407 (assemblage AII), respectively, were electroporated (GenePulser XCell, Bio-Rad) at 800 Ω, 350 V und 960 μF with ∼15µg linearized vector DNA. Transgenic parasites were selected in media containing 40 µg/ml puromycin and genome integration of the transgene was confirmed by PCR (not shown). Protein expression of ADI_AI_ transgene was assessed by Western blot on cell lysates as described before [63] using an anti-HA tag antibody (rat mab clone 3F10; Merck) and for loading control an anti-alpha-tubulin antibody (mouse mab clone DM1A; Merck). Peroxidase conjugated goat anti-mouse or anti-rat antibodies (1:5000, Bethyl laboratories) were used for detection (see above). To create the CRISPR-based ADI knockout, gRNAs were designed to target position 870 of the ADI gene (GL50803_112103) for insertion of antibiotic resistance cassettes. The gRNA (20 nucleotides) was designed to target the non-template strand (with 4-base overhangs complementary to the vector sequence overhangs using the Benchling software (https://benchling.com/crispr) with a NGG PAM sequence and the *G. duodenalis* ATCC 50803 genome (GenBank Assembly: GCA_000002435.1). gRNA oligonucleotides were annealed and cloned into Bbsl-digested Cas9U6g1pac as previously described [64]. Linear repair templates were designed with a left homology arm (750 bp upstream of the gRNA870 site), an antibiotic resistance cassette, and a right homology arm (750 bp downstream of the gRNA870 site). Two repair templates were designed: one with a blasticidin (bsd) resistance cassette and one with a hygromycin B (hyg) resistance cassette [42, 65]. Linear repair templates were synthesized by Twist Bioscience.

The CRISPR plasmid (Cas9U6gipac_112103g870) was electroporated into WBC6 trophozoites (20 μg DNA) as previously described [66]. The Cas9_112103g870 strain was maintained with antibiotic selection (50 μg/ml puromycin). The bsd linear repair template (20 µg) was first electroporated into the Cas9_112103g870 strain. Selection was maintained for the Cas9_112103g870 plasmid with 50 μg/ml puromycin and selection for integration of the repair template started at 75 μg/ml blasticidin, which was increased to 150 μg/ml blasticidin as the electroporated cells recovered. Blasticidin repair template integration was confirmed by PCR and long read sequencing. Next, the hyg linear repair template (20 µg) was electroporated into the Cas9_112103g870 strain that had the bsd repair template integrated at least once in the *Giardia* genome. Selection was maintained for the Cas9 plasmid with 50 μg/ml puromycin and selection for integration of the repair template started at 600 μg/ml hygromycin B, which was increased to 1200 μg/ml hygromycin B as the electroporated cells recovered. The hygromycin B repair template integration was confirmed by PCR and long read sequencing.

The KO strain confirmed to have both the bsd resistance cassette and hyg resistance cassette integrated into the genome with no wild type alleles was cloned by dilution to extinction. The contents of positive wells (those with growth) were transferred first to 6 ml TYI-S-33 medium, and then up to 12 ml TYI-S-33 medium. Clones were screened for complete knockout of the ADI gene by PCR (Supplemental Figure S6) using primers that bound to the 5’- and 3’-ends of the ADI gene (112103-F, 5’-AGATAGTCGTCGTGCACCTC -3’; and 112103-R, 5’-CAGATGTCAGCCTGCTCCAG-3‘). Upon insertion of an antibiotic cassette into the ADI gene, the PCR product would be increased by a known amount. One of the clones was selected for downstream analysis.

### ADI structure

For ADI of assemblage AI and assemblage AII isolates, the respective files from AlphaFold2 structure prediction [36] were downloaded and analyzed and illustrated using ChimeraX [67]. Also, Chai-1 [37] was used to predict arginine binding in the active center and to model structural changes by depicted amino acid mutations. The following datasets were used: AF-E2RU36-F1-model_v4 (WB6, assemblage AI) and AF-V6THM7-F1-model_v4 (DH, assemblage AII).

### Descriptive statistics

Data are given as mean ± standard deviations (SDs). Statistical significance was assessed by using paired (two-tailed) *t* test or one-way ANOVA with adequate post hoc tests, if not indicated otherwise. All analyses were performed with GraphPad Prism 9 software (GraphPad Software, Inc.) with a statistical significance level of *P* < 0.05.

### Data depository

ADI sequences presented are available at Genbank under following accession number: PV067229 - PV067249

## Acknowledgement and Funding information

We would like to thank Petra Gosten-Heinrich and Elke Radam for excellent technical lab assistance. The study was supported by internal grants of the Robert-Koch Institute (CK) and by the German Research Foundation (DFG) GRK2046 Research Training Programme (CK, TA) and Priority Programme SPP 2332 ‘Physics of Parasitism’ (CK, TA).

## Declaration of Interest

The authors declare no competing interests.

## Notes

### Competing Interest Statement

The authors have declared no competing interest.

