## Supplementary Figures for "Arginine metabolism has a pivotal function for the encystation of *Giardia duodenalis*"

**Supplementary Figure S1**: Representation of allelic sequences of ADI of 14 different patient isolates of *G. duodenalis* assemblage AII from an internal *G. duodenalis* biobank (Isolates: P029, P034, P064, P157, P168, P203, P212, P316, P324, P361, P368, P392, P407, P644). Sequences were retrieved by PCR and Sanger sequencing and aligned to reference ADI sequence “ADI_WB” of WB isolate (assemblage AI, gene ID “GL50803_112103”). Sequence of “P407” represents an identical sequence found in 11 different AII isolates. This sequence was also identical to the reference sequence of DH isolate (gene ID DHA2_112103). Three patient isolates had one or two additional mutations but at different sites in the sequences.

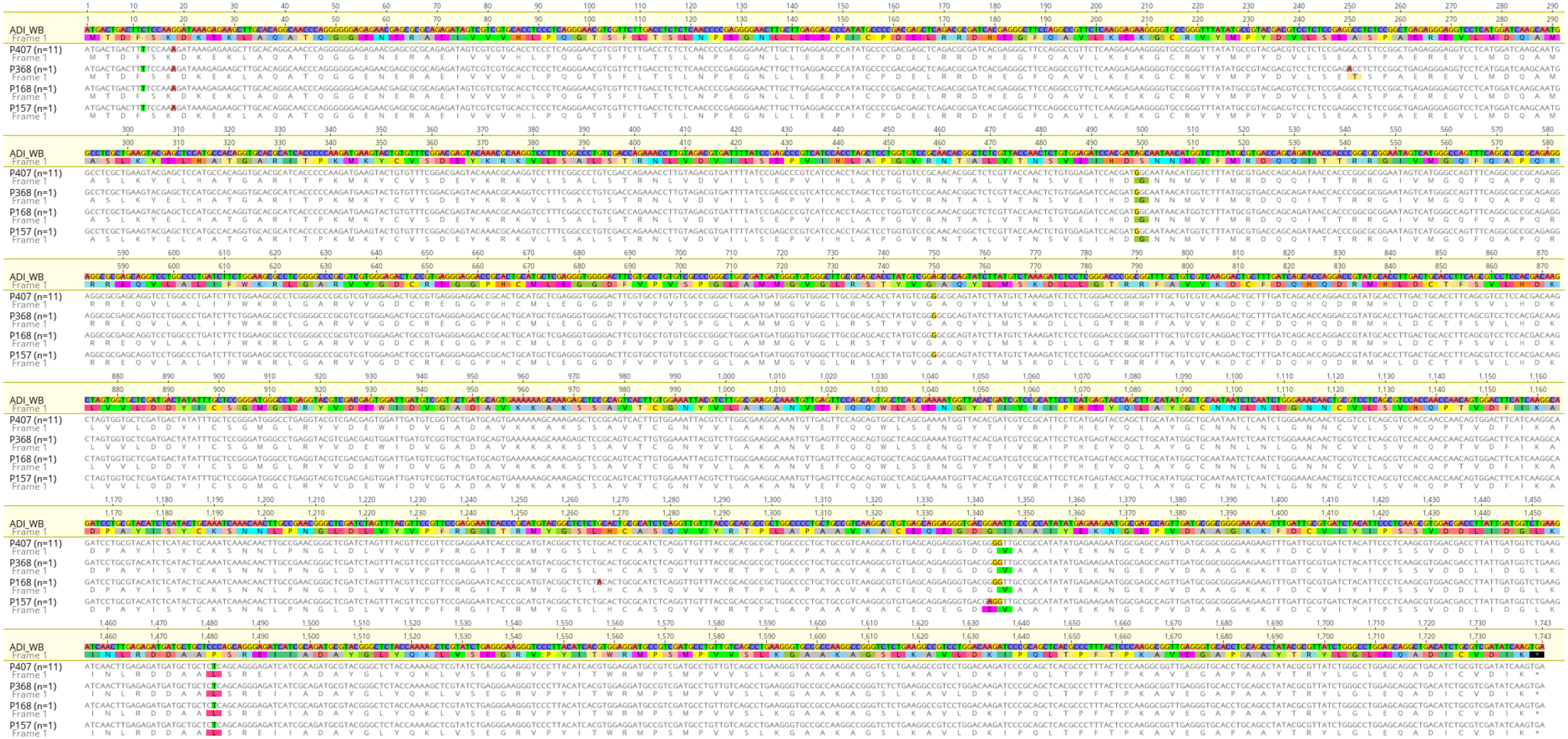

**Supplementary Figure S2**: Representation of allelic sequences of ADI of 15 different patient isolates of *G. duodenalis* assemblage B from an internal *G. duodenalis* biobank (Isolates: P132, P289, P344, P387, P413, P424, P427, P439, P448, P458, P486, P514, P621, P678, P786). PCR fragments of ADI of each isolate were cloned into pJet vector and transformed into E. coli. Plasmids of 10 single clones each were analyzed by Sanger sequencing and ADI sequences aligned to reference ADI sequence "ADI_GS" of GS isolate (assemblage B, gene ID "GL50581_1575"). Due to possible introduction of sequence chimeras during PCR, alleles were only defined "true" for sequences with two ore more identical copies withing the 10 analyzed clones. This revealed 17 different ADI alleles within the 15 isolates.

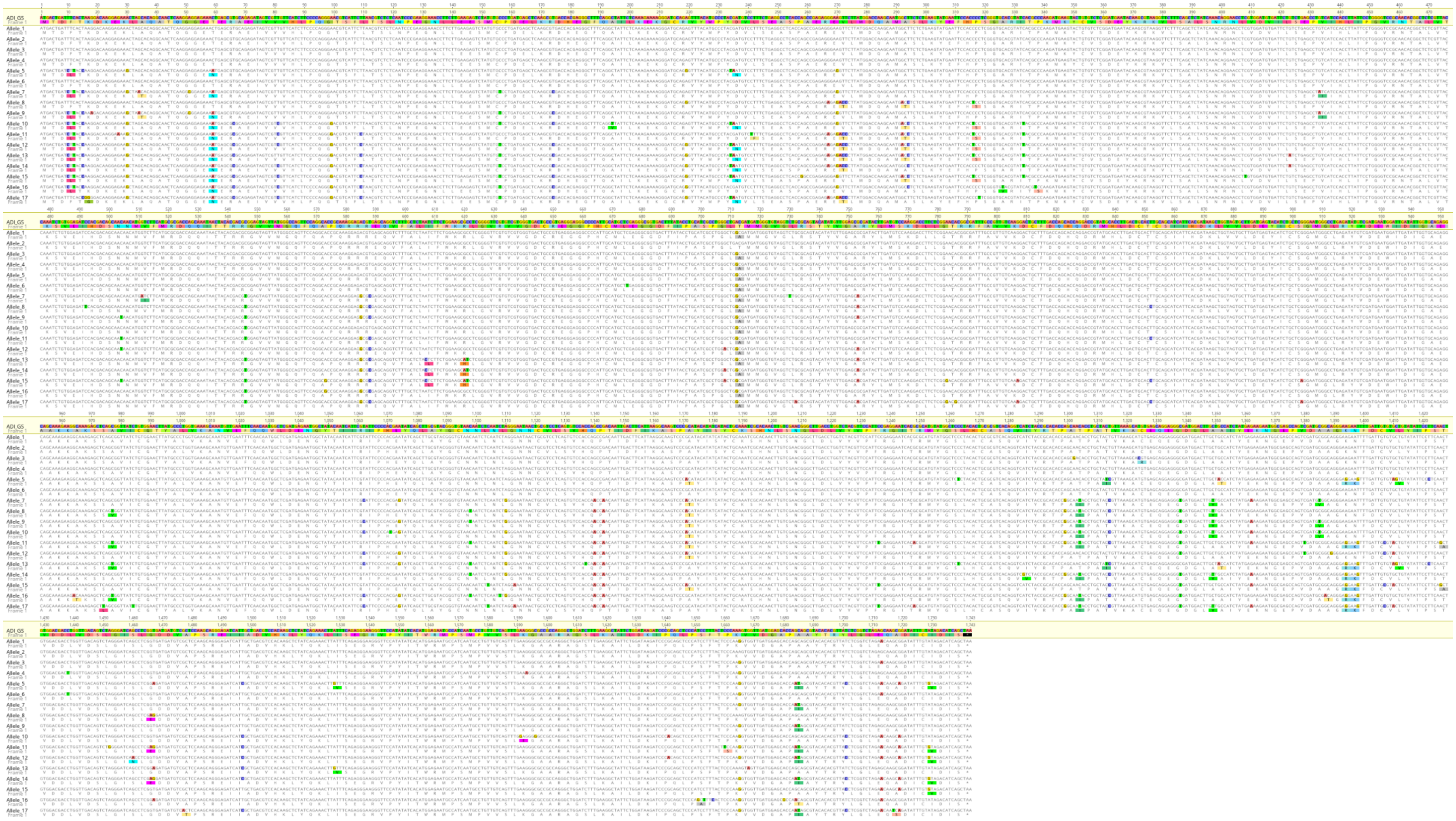

**Supplementary Figure S3:** Representation of all ADI protein variants derived from patient isolates of assemblage AII (see Fig S1) and assemblage B (see Fig S2) showing the conserved SNPs of assemblage AII sequences at position G167, V449 and L494. For comparison, reference sequences of assemblage AI (WB isolate) and assemblage B (GS isolate) were included. Note, AII_allele_1-4 represent corresponding sequenses of P407, P368, P168 and P157, respectively, shown in Supplementary Figure S1.

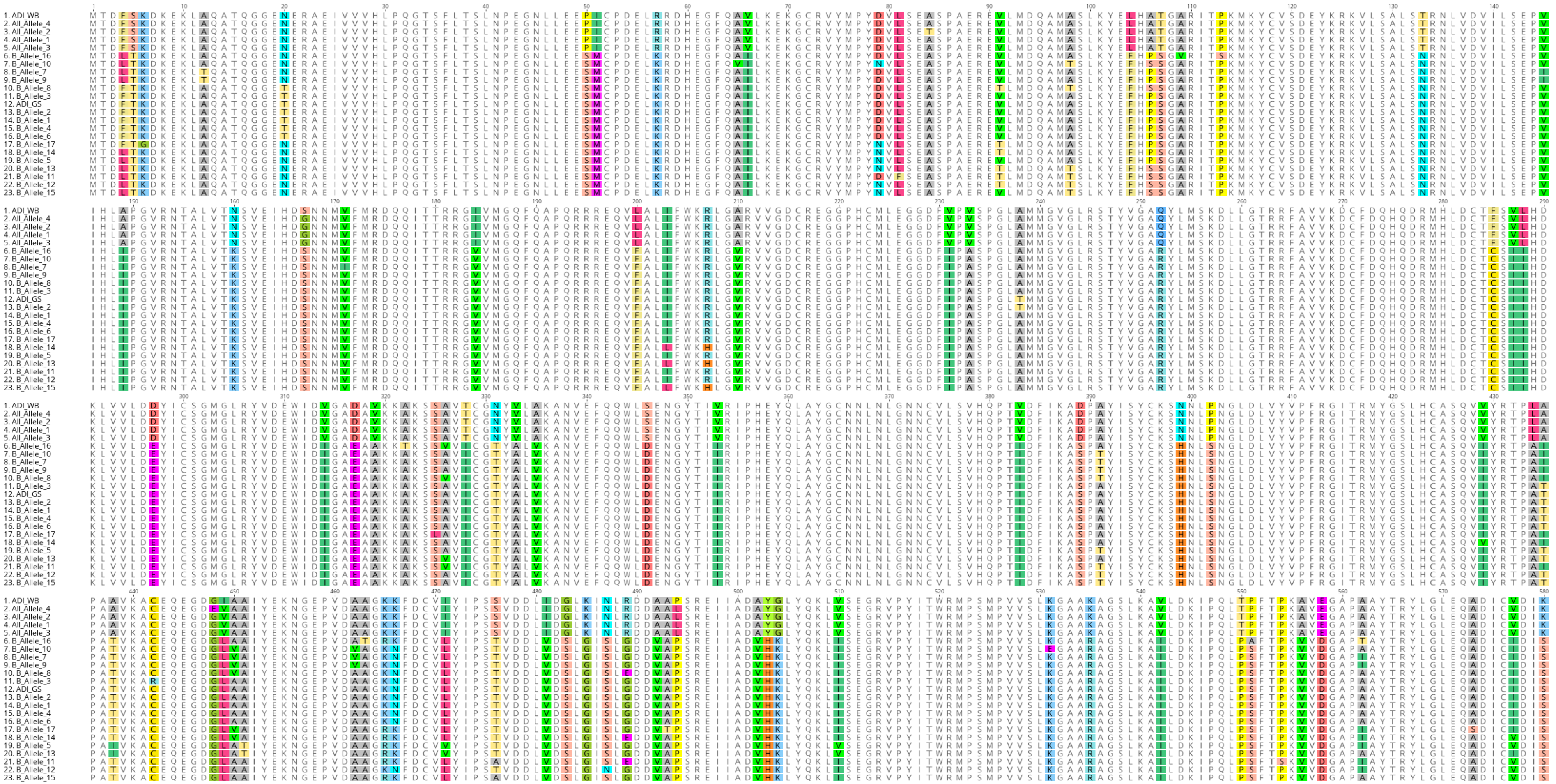

**Supplementary Figure S4**: Structural representation of $ADI_{AI}$ (Ser167, Ile449, Pro494) and overlay with mutated ADI functionally relevant in the assemblage AII ADI (Gly167, Leu494). Arginine substrate is positioned in the active center as predicted by the algorithm used in Chai-1 [1]. Color code is separated as indicated in the lower picture, RMSD (root mean square deviation) indicates the estimated difference in distance between the two modeled structures. pLDDT (predicted local distance difference test) represents reliability of the prediction.

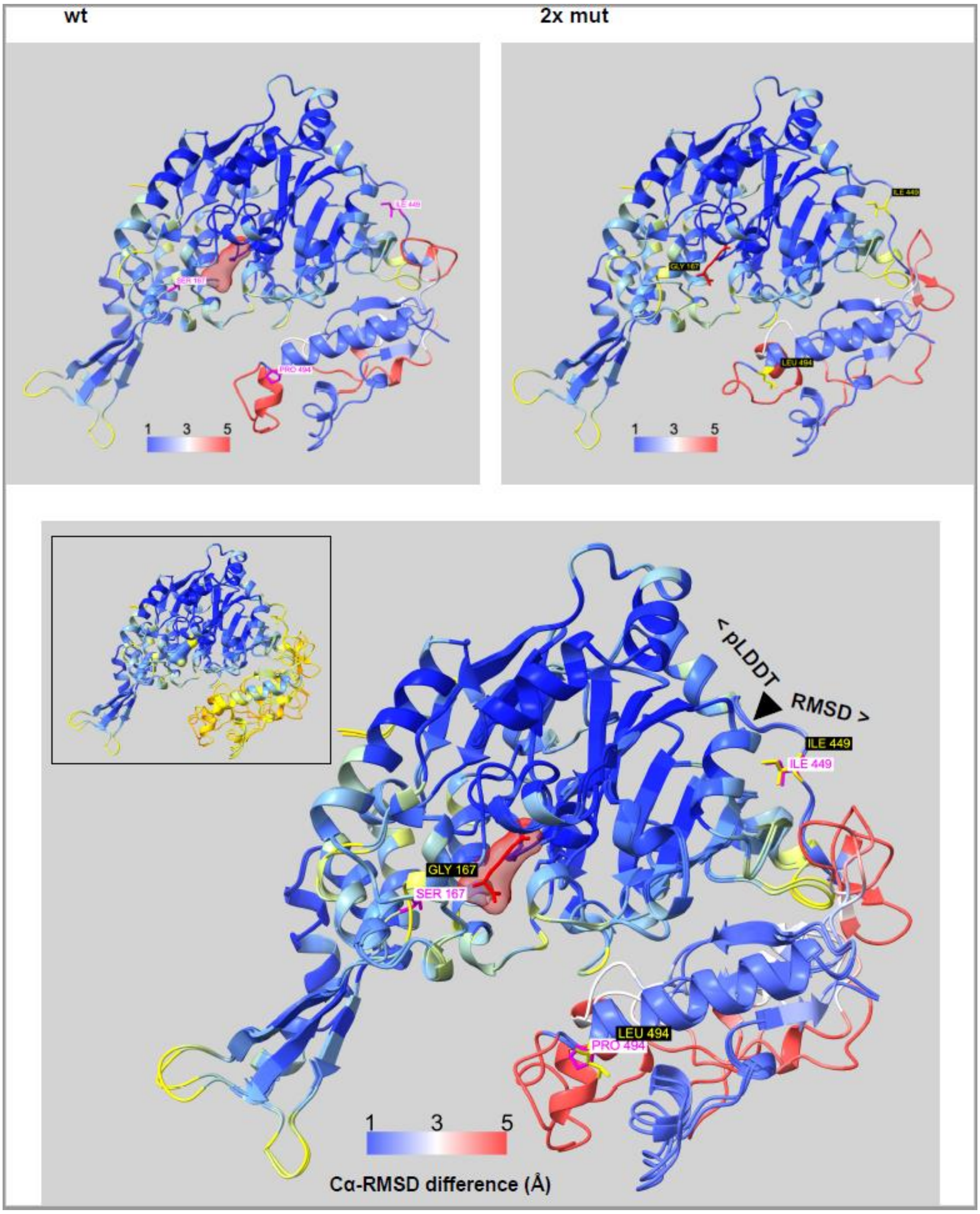

**Supplementary Figure S5:** (A) C57BL/6 mice infected with *Giardia duodenalis* assemblage B (GS/H7 strain, ATCC 50581) and fed with a defined arginine-free diet (n=14) showed significantly lower cyst excretion in the feces (pooled cyst numbers in feces collected day 4-7 post infection) than control animals (n=13) fed with the defined arginine-repleted diet containg 1% arginine. Note, that the defined diet is less rich and complex than "normal" diet, leading to overall lower cyst excretion. For comparison, mice fed with normal diet following the same infection protocol lead to significintly higher overall cyst excretion per gram feces at day 7 post infection ($5.1 \pm 1.3 \times 10^5$, n=22). (B) No differences were detected analyzing the genome equivalents by qPCR in tissue of the upper small intestine of the same mice as in (A). For statistics we used Kruskal-Wallis rank sum test.

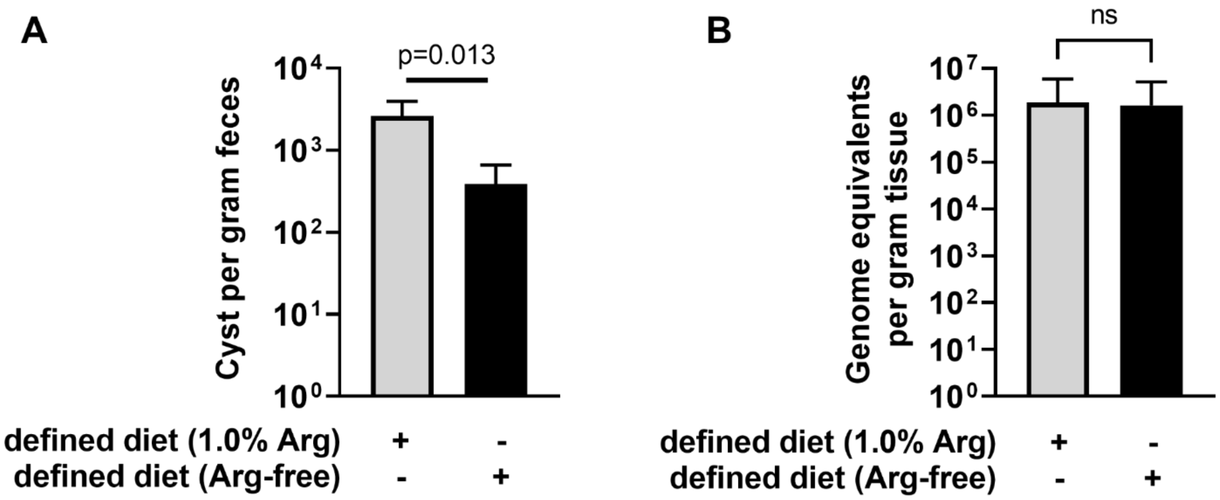

**Supplementary Figure S6:** PCR of ADI gene with primers flanking the antibiotics integration sites confirming ADIKO mutant. An ADI fragment was amplified that give rise to a size of ~1600 bp in the wildtype (WT, parental WB6 isolate), and of ~2450 bp (blasticidin resistance) and ~3075 bp (hygromycin resistance) in the ADIKO mutant. Control PCR without DNA is indicated by $H_2O$. PCR products were analyzed on a 1% agarose gel and vizualized using GelGreen reagent (Biotium, Fremont, CA) on a conventional documentation system (Vilbert Fusion FX6).

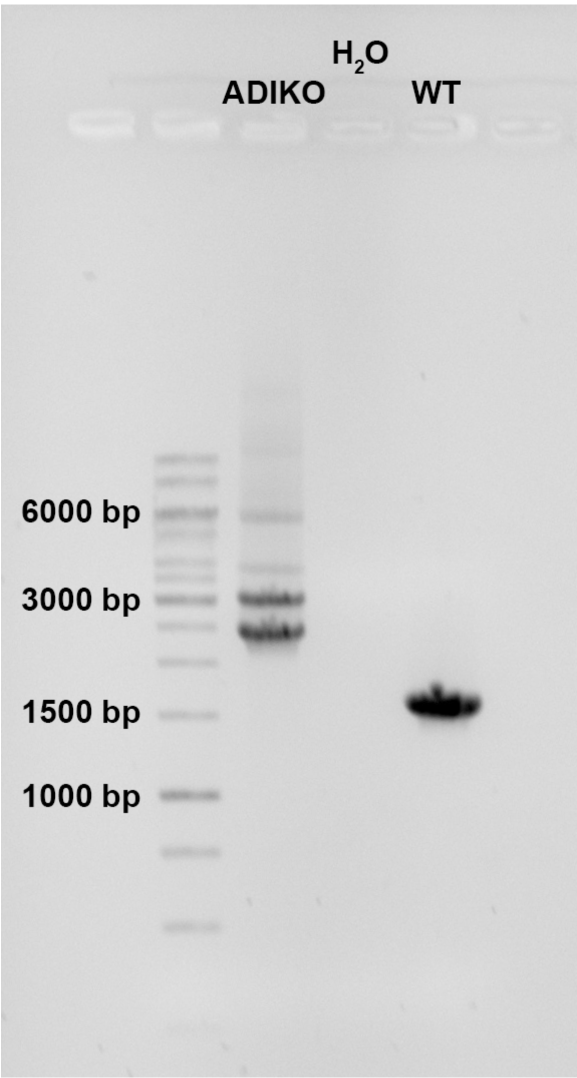
